# Agro-STARR-seq enables *in planta* genome-wide enhancer discovery and AI-empowerd enhancer design

**DOI:** 10.1101/2025.10.10.681754

**Authors:** Yongzhou Bao, Xiwen Zhang, Yanwei Song, Huiming Li, Junyan Qi, Hongqiu Cui, Na Li, Yun Chen, Weigang Zheng, Lingna Xu, Yuanyuan Zhang, Choulin Chen, Fangjie Zhu, Shaoqun Zhou, Zhaohong Li, Yuwen Liu

## Abstract

Comprehensive enhancer screening is essential for dissecting gene regulatory networks. However, methods for such screening remain limited in plants. We present Agro-STARR-seq, an *in planta* enhancer screening platform using *Agrobacterium*-mediated delivery of STARR-seq libraries into *Nicotiana benthamiana* leaves. In the *Arabidopsis* genome, we identified 14,715 sequences with enhancer activity, of which 91.59% were located outside of open chromatin regions. This largest *in planta* enhancer dataset to date enabled AI-assisted design of highly active synthetic enhancers. For the large *N. benthamiana* genome, we pre-processed the genomic DNA with an Assay for Transposase Accessible Chromatin (ATAC) procedure before the Agro-STARR-seq protocol,and found that only one-third of open chromatin exhibited enhancer activity, 76.00% of which showed no canonical H3K27ac histone marks. Footprinting analysis further enables systematic linkage of transcription factor (TF) binding to enhancer function. Overall, Agro-STARR-seq provides a novel platform for dissecting and engineering plant enhancers.

## Background

Gene expression governs plant cell fate determination, drives morphological development, and mediates adaptive responses to environmental stimuli [1]. This process is dynamically regulated by enhancers, which confer spatiotemporal precision and fine-tune transcriptional dynamics across developmental and environmental contexts [2]. Systematic identification of functional enhancers will facilitate the dissection of gene regulatory networks underlying complex traits and the *de novo* design of synthetic enhancers with desired activity.

Traditionally, plant enhancers have been inferred from epigenomic signatures, such as histone modifications and chromatin accessibility [3]. While these approaches are valuable, they rely on correlative biochemical markers and do not directly measure the enhancer activity. Methods based on DNA-binding activity of nuclear TFs were also developed [4], yet DNA-binding does not guarantee transcriptional-regulatory activity. The advent of massively parallel reporter assays (MPRAs) [5], exemplified by Self-Transcribing Active Regulatory Region Sequencing (STARR-seq) [6], enabled genome-wide, quantitative assessment of enhancer activity. Originally developed in *Drosophila melanogaster* [6] and *Homo sapiens* [7], STARR-seq has since been adapted to plants, including *Arabidopsis* [8], rice [9,10], maize [11,12] and barley [13], representing a major advance in functional plant genomics.

However, existing plant whole-genome STARR-seq studies have been limited to protoplasts (Supplemental Table S1). While protoplasts enable efficient delivery of enhancer-screening plasmid libraries—an essential requirement for STARR-seq—the enzymatic removal of cell walls and the loss of native tissue architecture could profoundly alter the intricate cellular states transcriptional regulatory networks in the multicellular context. Consequently, enhancers identified in protoplasts may not accurately reflect their activity in intact tissues and may fail to capture many genuine enhancers, leaving a major gap in our understanding of plant gene regulation *in planta*. To date, genome-wide STARR-seq enhancer profiling directly *in planta* has not been achieved.

To address this limitation, we developed Agro-STARR-seq (*Agrobacterium*-mediated Self-Transcribing Active Regulatory Region Sequencing, Fig. 1A), a technology that enables high-throughput, genome-wide profiling of enhancer activity directly in plant leaves. This approach preserves the native regulatory environment while providing quantitative assessment of enhancer strength. Applying Agro-STARR-seq to the entire *Arabidopsis thaliana* genome, we demonstrated its capacity for comprehensive enhancer discovery and highlighted the utility of the resulting dataset for AI-empowered *de novo* design of synthetic enhancers that surpass the activity of natural sequences [14]. We further applied Agro-STARR-seq to systematically identify enhancers within the open chromatin regions of *Nicotiana benthamiana* leaves, providing new insights into the epigenomic features and the *cis*-regulatory lexicon of plant enhancers.

**Figure 1.**
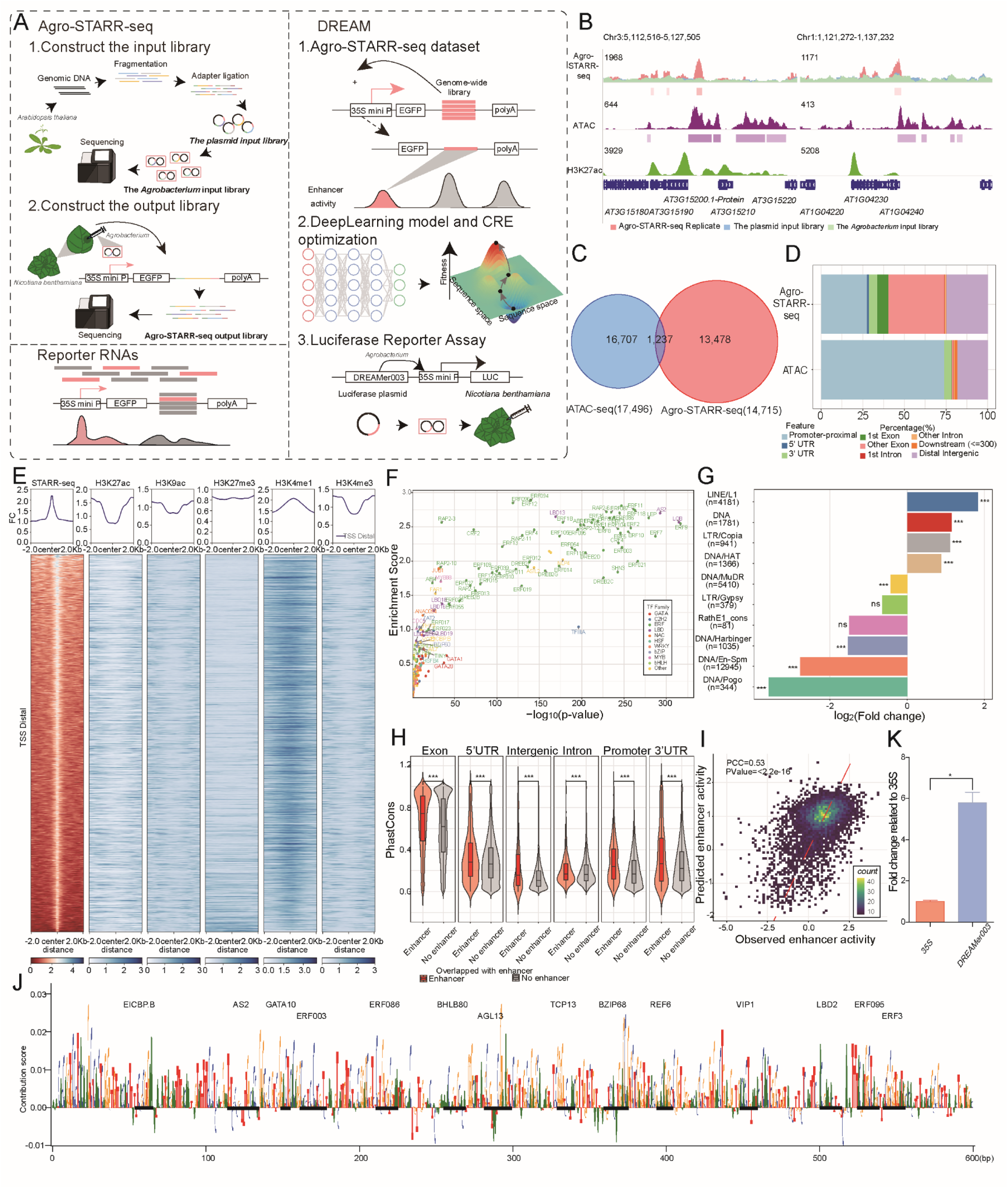
*In planta* Identification and Engineering of *Arabidopsis* Enhancers Using Agro-STARR-seq and DREAM Framework. (A) Schematic overview of Agro-STARR-seq workflow and DREAM framework. Agro-STARR-seq generates high quality *in planta* enhancer data which is subsequently used as training data by DREAM to design synthetic enhancers. Agro-STARR-seq input libraries include the plasmid input library and *Agrobacterium* input library. The plasmid input library: Genomic DNA is fragmented and cloned into a modified Agro-STARR-seq vector containing a 35S mini promoter driving basal EGFP expression. The *Agrobacterium* input library: The plasmid input library is transformed into *Agrobacterium* by electroporation. Agro-STARR-seq output library: The *Agrobacterium* input library is infiltrated into *N. benthamiana* leaves. After 72 hours, RNA is isolated for next-generation sequencing. Enhancer activity is quantified by comparing sequence enrichment in the output library to the input library. DREAM comprises two integral modules: the state-of-the-art SENet that predicts enhancer activity using DNA sequences as the input and the evolutionary optimization module that designs strong synthetic enhancers by employing a genetic algorithm, which iteratively maximizes regulatory activity as predicted by the SENet-derived model. (B) Representative genomic regions show the signal profiles from both output and input libraries, alongside ATAC-seq and H3K27ac tracks. The first genome browser track displays fragment densities from the Agro-STARR-seq output library (red), the plasmid input library (blue), and the *Agrobacterium* input library (green). (C) Venn diagram depicting overlap between Agro-STARR-seq enhancers and ATAC-seq peaks [16]. (D) Annotation of regulatory regions relative to the transcriptional start site (TSS) (UTR: untranslated region; Promoter-proximal: ±500 bp TSS-proximal regions; Distal intergenic: all regions outside the promoter-proximal, gene body, and 300 bp downstream regions). (E) Heatmaps show signal enrichment (±2 kb from enhancer centers) for STARR-seq, H3K27ac, H3K9ac, H3K27me3, H3K4me1, and H3K4me3 marks, categorized by TSS proximity, with the figure highlighting TSS distal regions. Signal intensity represents fold enrichment of ChIP-seq over input (normalized by library size, detailed in Supplemental notes). (F) TFs motifs enriched in the Agro-STARR-Seq enhancers compared to genomic regions of non-enhancers (background). The x -axis represents the -log_10_p-value, and the y -axis represents the Enrichment score. (G) Enrichment profile of enhancers overlapping with TEs. (H) Average PhastCons scores of Agro-STARR-seq enhancers and non-enhancers in different genomic categories. (I) Performance evaluation of DREAM enhancer activity prediction module using hold-out chromosome dataset from *A. thaliana*. (J) Nucleotide contribution scores for the optimized enhancers derived from the enhancer activity models using DeepExplainer. Instances of motifs identified by DREAM are emphasized, with known motifs indicated in black. (K) Luciferase reporter assay validation of a DREAM-optimized synthetic enhancer. Values normalized to Renilla luciferase signal. Error bars: Standard error of the mean (n = 3 biological replicates). This synthetic enhancer was named DREAMer003, in line with the strong enhancer DREAMer001 and silencer DREAMer002 designed using Drosophila STARR-seq data in our previous study [14].

## Results and discussion

### Development and Validation of Agro-STARR-seq System

The Agro-STARR-seq method exploits *Agrobacteria tumefaciens* to deliver STARR-Seq enhancer screening plasmids into intact leaves of *Nicotiana benthamiana.* To enable this, we modified the conventional STARR-seq vector for compatibility with *Agrobacterium*-mediated delivery (Fig. S1A and S1B, detailed in Supplemental notes). As a proof of concept, we constructed an enhancer-screening library from randomly fragmented *Arabidopsis thaliana* Col-0 genomic DNA with an average fragment size of 450 bp, taking advantage of its relatively small genome size (ca. 135 Mbp) to streamline experimental procedures (Fig. S2A). To evaluate library complexity, we sequenced both the plasmid input library and the corresponding *Agrobacterium* input library after transformation. The latter contained at least 4.8 million independent candidate fragments (Supplemental Table S2), achieving 94% coverage of the *A. thaliana* Col-0 genome (Fig. S2B). Simulation-based analysis further indicated that the library complexity had reached saturation (Fig. S2C). Fragment abundance between plasmid and *Agrobacterium* input libraries showed high correlation (Pearson’s correlation coefficient, PCC = 0.9, P-value < 2.2e^−16^, Fig. S3A), confirming minimal sequence bias during transformation. After infiltrating *N. benthamiana* leaves with the screening library–containing *Agrobacteria*, transcript abundance from STARR-seq plasmids showed strong reproducibility across two biological replicates (PCC = 0.92, P-value < 2.2e^−16^, Figure S3B). Collectively, these results demonstrate that Agro-STARR-seq achieves library complexity and coverage comparable to established STARR-seq protocols published from our previous work [7,15], validating the robustness of this method.

### Genome-wide Enhancer Atlas of the *Arabidopsis* Genome

To quantitatively assess enhancer activity, we compared the enrichment of cDNA fragments of the output libraries relative to *Agrobacterium* input library DNA fragments across genomic bins. Sequences exhibiting increased self-transcription are indicative of stronger enhancer activity (Fig. 1B and S4A-G). Using this approach, we identified 14,715 candidate enhancers *in planta*, with activities highly reproducible across replicates. Notably, only 1,237 (8.4%) of these enhancers were located within previously annotated open chromatin regions (Fig. 1C) [16], consistent with previous *in vitro* studies in rice [11] and human [7]. This finding suggests that a substantial proportion of sequences with *in planta* enhancer potential reside outside open chromatin regions.

We next characterized the genetic and epigenetic features of the enhancers. *In planta A. thaliana* enhancers (mean peak length = 200 bp; Fig. S5A) were distributed across all five *A. thaliana* chromosomes, with certain genomic hotspots exhibiting higher densities (Fig. S5B and S5C). Genomic annotation revealed that 27.5% of identified enhancers were located promoter-proximal regions, while 24.8% resided in distal intergenic regions (Fig. 1D). This preferential distribution near genes likely reflects the compact genome architecture of *A. thaliana*. Integration with ChIP-seq profiles for five histone modifications (H3K27ac, H3K9ac, H3K27me3, H3K4me1, and H3K4me3; Supplemental Notes) showed that promoter-proximal enhancers co-localized with H3K4me3, H3K27ac, and H3K9ac peaks, while distal enhancers were associated with H3K4me1 (Fig. 1E and S6), consistent with previously reported ChIA-PET results for *A. thaliana* [16].

Motif analysis revealed 108 significantly enriched transcription factors (TFs) DNA binding motifs (q-value < 0.05), including members of the ERF family, which are widely involved in the regulation of plant growth, development, and environmental responses (Fig. 1F). Many enhancers co-localized with transposable elements (TEs), predominantly with LINE/L1 (4,181), LTR/Copia class I retrotransposons (941), and class II transposons including DNA (1,781) and DNA/HAT (1,366) (One-sided hypergeometic test; Fig. 1G), indicating specific TE types may contribute significantly to enhancer formation in *A. thaliana*. Furthermore, a pronounced differential enrichment of class II DNA transposons (1,781) between active and inactive accessible chromatin regions was observed, suggesting that specific TEs contribute differently to enhancer activity and chromatin accessibility (Fig. S7). Evolutionary rate analysis demonstrated these enhancers exhibit significantly higher conservation than non-enhancers across different functional genomics categories (Fig. 1H). Collectively, these results highlight the functional and biological significance of the enhancers identified through our Agro-STARR-seq approach *in planta*.

### AI-empowered *de novo* plant enhancer design from *in planta* activity

Unlike previous protoplast-based plant STARR-seq studies, this proof-of-concept application of Agro-STARR-seq scanned the entire *A. thaliana* genome, generating the largest *in planta* enhancer dataset to date—an invaluable resource for AI-driven enhancer design in plant leaves. Using this dataset, we applied our recently developed computational framework, DREAM (DNA *cis*-Regulatory Elements with Controllable Activity Design PlatforM) (Fig. 1A) [14] to design synthetic enhancers with high regulatory activity in *N. benthamiana* leaves. The enhancer prediction module of DREAM demonstrated robust generalization capability, achieving a PCC of 0.53 (P-value < 2.2e^-16^, Fig. 1I, detailed in Supplemental notes) between predicted and observed enhancer activities in a hold-out chromosome dataset. Its sequence design module uses a genetic algorithm to iteratively maximize DREAM-predicted enhancer activity, generating synthetic enhancers predicted to exhibit exceptionally high regulatory activity. Simulated evolution analysis showed that enhancer activity reached saturation after approximately 100 generations (Fig. S8). From the 100th generation, we randomly selected a synthetic enhancer which contained several key TF motifs, including ERF003 and ERF095, known activators of gene expression. A model interpretability analysis confirmed the substantial contribution of TFs motifs to enhancer activity prediction (Fig. 1J). Furthermore, *in planta* luciferase reporter assays (detailed experimental methods are provided in the Supplemental notes, with vector constructs shown in Fig. S9A and S9B) validated the functionality of this DREAM-designed synthetic enhancer (DREAMer003), demonstrating a 5.8-fold activity increase when positioned upstream of a viral 35S promoter (One-sided t-test, P-value < 0.05, Fig. 1K and Supplemental Table S3).

### ATAC-Agro-STARR-seq identifies enhancers of the *N. benthamiana* genome

To demonstrate the versatility of our Agro-STARR-seq platform beyond *A. thaliana*, we extended its application to identify enhancers in open chromatin regions of *N. benthamiana* leaves (ATAC-Agro-STARR-seq, Fig. 2A). Notably, this setup provides a cellular environment fully matched to the genomic sequences under investigation, offering a robust framework to dissect the genetic, epigenetic, and functional properties of plant enhancers. To enrich for open chromatin regions in the enhancer screening vector, we adapted the ATAC-seq protocol and generated the first high-quality ATAC-seq library from *N. benthamiana* leaves (Supplemental Table S4) [17], comprising ∼88 million fragments largely representing nucleosome-free regions (NFRs, Fig. S10A and S10B). This library displayed hallmark epigenetic features, including strong enrichment at transcription start sites (TSSs, Fig. S10C), peak sizes of 250–750 bp (Fig. S10D), and predominant localization in distal intergenic and promoter regions (Fig. 2B). Functionally, genes proximal to ATAC-seq peaks exhibited significantly higher expression than distal genes (Fig. S10E and S10F), confirming the relevance of these accessible regions.

**Figure 2.**
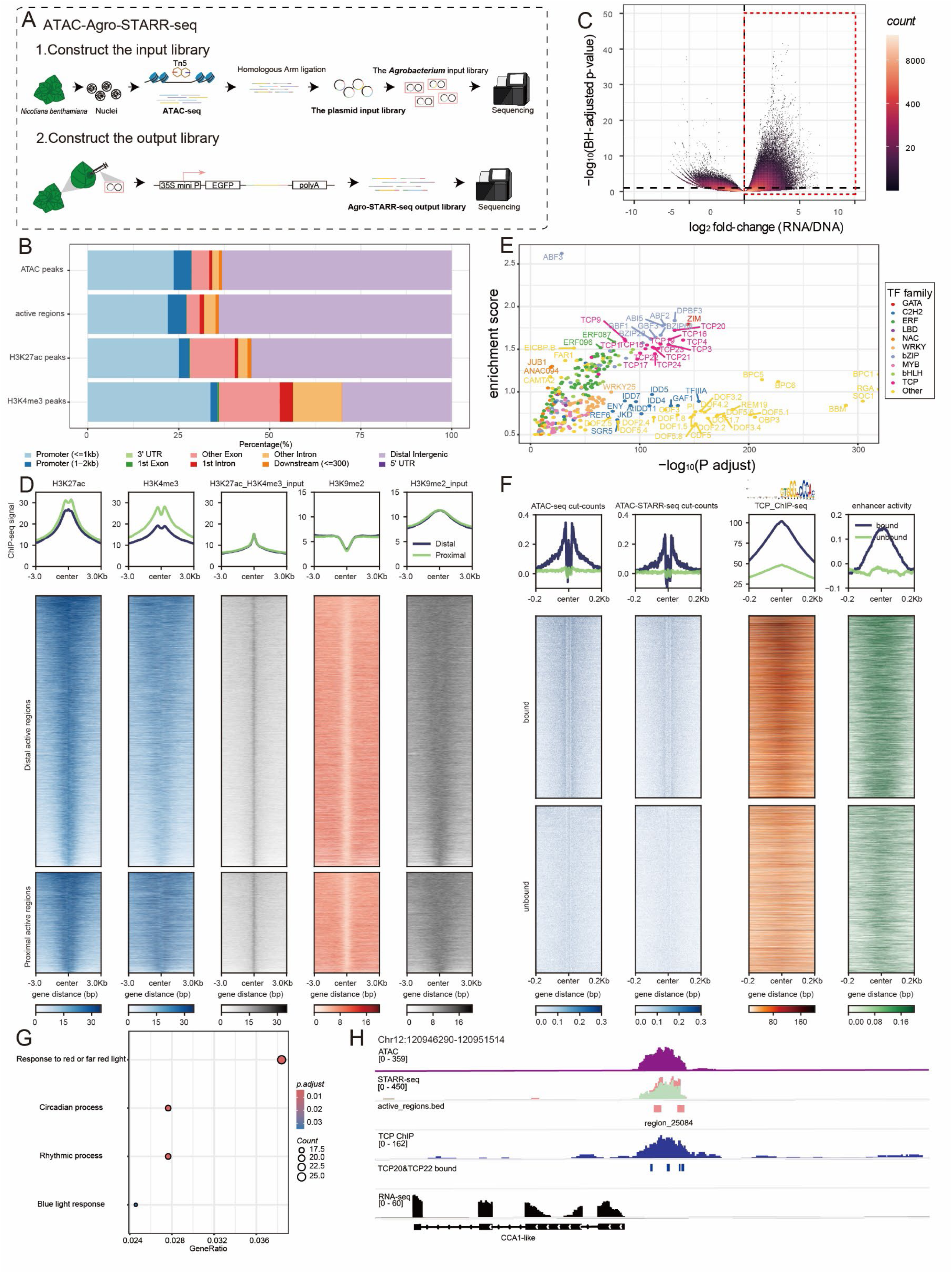
*In planta* enhancer screening and characterization of *N. benthamiana* leaf open chromatin regions using ATAC-Agro-STARR-seq. (A) Schematic overview of ATAC-Agro-STARR-seq workflow. ATAC-Agro-STARR-seq input libraries include the plasmid input library and *Agrobacterium* input library. The plasmid input library: Open chromatin is isolated from *N. benthamiana* leaf nuclei with the cut-and-paste transposase Tn5. The open chromatin fragments are cloned into the Agro-STARR-seq plasmid. The *Agrobacterium* input library: The plasmid input library is transformed into *Agrobacterium* by electroporation. Agro-STARR-seq output library: The *Agrobacterium* input library is infiltrated into *N. benthamiana* leaves. After 72 hours, RNA is isolated for next-generation sequencing. Enhancer activity is quantified by comparing sequence enrichment in the output library to the input library. (B) Annotation of open chromatin peaks, their active regions (Agro-STARR-seq enhancer in open chromatin regions), and H3K27ac and H3K4me3 peaks relative to the transcription start site (TSS). Promoters were defined as regions extending 2 kb upstream and 1 kb downstream of the TSS. (C) A scatter plot showing the log₂ fold change (log_2_FC) of normalized output to input coverage, and the -log_10_ (BH-adjusted p-value) for all bins, as identified by DESeq2 analysis. Red boxes indicate active bins. (D) Heatmaps of histone modification ChIP-seq signals in ATAC-Agro-STARR-seq-defined active regulatory elements (proximal and distal). The columns (left to right) are as follows: 1-2) H3K27ac and H3K4me3 signals; 3) the corresponding input control for the active marks; 4) H3K9me2 signal, a heterochromatin mark shown for comparison; 5) input control for H3K9me2. (E) Enrichment analysis of transcription factor motifs in ATAC-Agro-STARR-Seq enhancer regions. (F) Comparison of ATAC-STARR-seq cut count signal, ATAC-seq cut count signal, TCP ChIP-seq signal, and enhancer activity between predicted motif-bound and predicted motif-unbound regions of TCP20. G) GO enrichment analysis of nearest-neighbor gene sets associated with TCP23 bound active accessible peaks (Supplemental Table S10). H) Chromatin accessibility, TCP binding sites, TCP ChIP signals, active regions, and RNA-seq profiles at the CCA1-like gene.

Cloning this high-quality ATAC-seq library into the Agro-STARR-seq vector produced highly consistent fragment coverages across subsequent plasmid input libraries and *Agrobacterium* input libraries, with an average complexity exceeding 10 million unique fragments (Fig. S11 and S12; Supplemental Table S3). We then infiltrated the ATAC-Agro-STARR-seq *Agrobacterium* input libraries into *N. benthamiana* leaves, generating output libraries and identifying 39,394 enhancers in open chromatin regions (hereafter termed active accessible regions, Fig. 2C). Notably, only ∼33.2% of accessible chromatin peaks contained at least one active enhancer, highlighting the limitations of using ATAC-seq signal alone when annotating plant enhancers [18]. The genomic distribution of active accessible regions closely mirrored that of ATAC-seq peaks, indicating no strong positional bias within open chromatin (Fig. 2B). Active accessible regions were associated with higher expression of neighboring genes, demonstrating their functional relevance in the native chromatin context (Fig. S13A).

Interrogation of published histone modification datasets (H3K27ac, H3K4me3, H3K27me3, and H3K9me2) revealed that these active regions were significantly enriched for marks of active chromatin (H3K27ac, H3K4me3) but not the repressive (H3K27me3) or heterochromatin (H3K9me2)-associated mark (Fig. S13B; Extended Data Table). We further observed distinct patterns of histone mark enrichment at proximal versus distal sites: while the promoter-associated H3K4me3 mark exhibited higher signals specifically in proximal active regions, H3K27ac was enriched at both proximal and distal active regions (Fig. 2D).

Notably, we found that 76.00% of enhancers identified within open chromatin regions lacked H3K27ac, a canonical enhancer-associated histone modification, revealing tens of thousands of previously hidden enhancers in the plant genome. This is consistent with previous studies in human and mouse [19], and underscores the limitation of relying solely on active histone marks to annotate plant enhancers.

### Integration of accessibility and activity reveals TF binding and functional networks

To identify TFs contributing to enhancer activity, we analyzed TF DNA binding motifs enriched in all, proximal, and distal active accessible regions. Many TFs, including members of the TCP, ERF, bZIP, and bHLH families, were significantly enriched in these regions (Fig. 2E). Notably, TCPs, previously reported as major contributors to distal chromatin accessibility quantitative trait loci (caQTLs) in maize, were most significantly enriched in distal active accessible regions (Fig. S14) [12]. These results demonstrate that our ATAC-Agro-STARR-seq platform provides a robust framework for providing insights into how TFs contribute to plant enhancer function.

To further link TF binding with enhancer activity, we integrated the footprinting-based TF binding site predictions from ATAC-seq with the precise enhancer activity measurements of STARR-seq. By combining the strengths of both approaches, Agro-STARR-seq offers a unique platform to systematically investigate the relationship between TF binding and enhancer activity, providing an integral perspective to study concerted *cis-trans* transcriptional regulation in plants.

To validate this approach, we generated Tn5-bias–corrected cut site signal files from ATAC-Agro-STARR-seq data and predicted TF binding sites using footprint scores. TFs with available ChIP-seq data in *N. benthamiana* leaves served as a gold standard for evaluating predicted versus actual binding sites. For example, for TCP20, corrected cut site signals clearly distinguished predicted TF-bound from unbound sites (Fig. 2F and S15), a trend consistent across other TFs (Fig. S16). ChIP-seq confirmed preferential binding of TCP20 to the predicted sites, and integration with STARR-seq data demonstrated significantly higher enhancer activity at these bound sites compared to unbound sites, providing genome-wide evidence that TCP20 binding generally activates gene expression.

Subsequently, we focused on the TCP regulatory network and examined the functions of genes associated with predicted TCP-bound active accessible peaks. We found that TCPs regulate genes involved in circadian rhythm and light responses (Fig. 2G and Fig. S17). TCPs have been reported to bind directly upstream of the CCA1 transcription start site, a morning gene in *A. thaliana*, and activate its expression [20,21]. In *N. benthamiana*, CCA1 also functions in circadian rhythm regulation [22]. Although CCA1 is not annotated in the current *N. benthamiana* reference genome (Niben261), we identified and assembled a CCA1-like transcript from RNA-Seq data, mapped to Niben261 Chr12:120,946,470–120,948,272 (Supplemental Table S5 and S6). Strikingly, we found that near the promoter of this transcript, the predicted TCP-bound sites upstream of this putative NbCCA1 (as supported by published TCP ChIP-seq signals, also exhibit Agro-STARR-Seq enhancer activity (Fig. 2H).

Collectively, our findings establish *in planta* ATAC-Agro-STARR-seq as a powerful method for delineating regulatory networks, as it directly connects transcriptional activation by upstream TFs to the expression of their downstream target genes.

## Conclusions

Here, we present Agro-STARR-seq, a high-throughput method that addresses the challenge of transitioning high-throughput enhancer screening from isolated protoplast systems to living plants. The approach utilizes *Agrobacterium*-mediated delivery into intact *N. benthamiana* leaves—a widely adopted model plant that provides a rapid, scalable, and transient gene expression platform, thus establishing an ideal *in planta* chassis. Whereas previous *in planta* methods screened only thousands of sequences for enhancer activity (Supplemental Table S1) [23], Agro-STARR-seq scales this up by more than 100-fold, enabling genome-wide enhancer discovery across diverse plant species. Beyond uncovering novel genetic and epigenetic features of plant enhancers, Agro-STARR-seq generates large-scale *in planta* enhancer datasets suitable for AI training, offering a unique platform for precise engineering of transcriptional regulatory networks in plant genomes.

For method development, we first screened enhancer activity of the entire *A. thaliana* genome. *A. thaliana* was chosen as the source of test sequences due to its small genome size, which facilitates methodological development and efficient validation. Nonetheless, the substantial increase in the scale of our methodology enabled a transition from assessing enhancer activities in a limited set of candidates to broader genomic regions or genomic regions enriched through functional genomics techniques such as ATAC-seq and CUT&Tag. This versatility establishes Agro-STARR-seq as a broadly applicable platform for functional annotation of plant enhancers.

To further explore the genetic and epigenetic features of plant enhancers, we profiled enhancer activity in the accessible chromatin regions in *N. benthamiana* by ATAC-Agro-STARR-Seq. In this study, we successfully constructed ATAC-seq libraries in *N. benthamiana* for the first time, identifying 99,851 peaks with a FRiP (Fraction of Reads in Peaks) score of 34.90% (Supplemental Table S4). Integration of footprinting signals of ATAC-seq and Agro-STARR-seq enables the simultaneous capture of multiple regulatory layers—including chromatin accessibility, transcription factor binding, and enhancer activity—within a single assay. As demonstrated in *N. benthamiana*, this integrated profiling offers a powerful strategy to unravel multilayered gene regulatory networks, exemplified by our discovery of a TCP-mediated circadian regulatory circuit. Strikingly, we found that 76.00% of enhancers in *N. benthamiana* leaf accessible chromatin regions lack H3K27ac marks, revealing—for the first time in plants—the widespread presence of ’hidden’ enhancers, similar to observations in animals [19]. This highlights the complexity of inferring enhancer activity from histone modifications and underscores the value of Agro-STARR-seq in plant genetics. Current large-scale, histone mark-based enhancer screens may overlook many functionally important enhancers and enhancer-modulating variants that shape plant phenotypes.

The successful integration of our high-throughput screening platform with the DREAM computational framework marks a paradigm shift in plant synthetic biology. By combining the largest *in planta* enhancer activity dataset to date with deep learning– driven design, we engineered synthetic enhancers exhibiting a 5.8-fold increase in regulatory activity over the conventional viral 35S promoter, long a benchmark for transgenic expression. This unprecedented enhancement demonstrates the power of bridging large-scale functional genomics with predictive AI models to surpass the limitations of natural cis-regulatory elements. Since N. *<u>benthamiana</u>* responds to multiple stresses, such as heat and drought, generating Agro-STARR-seq data under diverse conditions will enable AI-guided design of environment-responsive enhancers—crucial for developing ‘smart’ plants that fine-tune critical gene expression to adapt to environmental challenges.

In conclusion, our work opens new avenues for deciphering gene regulatory networks and accelerates the engineering of tailored traits in plants, marking a major step forward in plant functional genomics and plant synthetic biology.

## Methods

### Construction of Agro-STARR-Seq enhancer screening vector

To construct a vector suitable for Agro-STARR-seq experiments, we modified the reporter gene vector based on the backbone of the pGreen Ⅱ plasmid (Supplemental Table S7, Fig. S1). In brief, the CaMV 35S promoter was replaced with the CMV 35S minimal promoter, and the CaMV poly (A) signal was removed. Additionally, several key functional elements were incorporated, including an intron, EGFP sequence, terminator elements, and a CmR-ccdB cassette. These sequences were arranged sequentially and are underlined in Fig. S1.

### Preprocessing of *Arabidopsis* whole-genome plasmid input Library

Three micrograms of *Arabidopsis thaliana* Col-0 genomic DNA was diluted to 10 ng/μL and fragmented using ultrasonication (Diagenode). The fragmented DNA was size-selected (450–500 bp) through a 1% agarose gel (D4008; ZYMO). DNA fragments underwent end repair, adapter ligation, and purification (ND607; Vazyme Biotech) (N411; Vazyme Biotech). Adapter-ligated DNA was then amplified using polymerase chain reaction (PCR) with the KAPA HiFi HotStart ReadyMix PCR Kit (KK2602; Roche), with homologous arms incorporated for cloning (Supplemental Table S8). The PCR cycling conditions were: 98°C for 45 s; 15 cycles of 98°C for 15 s, 65°C for 30 s, and 72°C for 70 s. Amplified products were purified and used for downstream cloning into the Agro-STARR-seq vector (T1030S; NEB).

### Preprocessing of *N. benthamiana* ATAC-seq plasmid input library

Leaves (500 g) were harvested from two vigorously growing *Nicotiana benthamiana* plants (approximately one month old, pre-flowering stage). Nuclei were isolated using the Plant Sample Nuclei Isolation Kit (52305-10, BioYou) according to the manufacturer’s protocol. Assay for transposase-accessible chromatin (ATAC) was conducted using Chromatin Profile Kit (N248-01A, Novoprotein) following the manufacturer’s instructions, respectively.

*N. benthamiana* ATAC-seq library was then amplified using polymerase chain reaction (PCR) with the KAPA HiFi HotStart ReadyMix PCR Kit (KK2602; Roche), with homologous arms incorporated for cloning (Supplemental Table S8). The PCR cycling conditions were: 98°C for 45 s; 15 cycles of 98°C for 15 s, 65°C for 30 s, and 72°C for 70 s. Amplified products were purified and used for downstream cloning into the Agro-STARR-seq vector (T1030S; NEB).

### Construction of the plasmid input libraries

The *A. thaliana* and *N. benthamiana* purified PCR products, now containing homologous arms, were inserted into the linearized vector backbone (digested with AgeI and SacI) via Gibson assembly (E5510S; NEB). After recombination, all products were pooled, and 5 μL of DNA constructs were subsequently transformed into MegaX DH10B™ T1R electrocompetent cells (C640003; ThermoFisher Scientific). A total of four Gibson assembly and transformation experiments were performed. All transformed Escherichia coli cultures were pooled and grown in 1 L of LB medium until the optical density (OD) reached 1.0. The plasmid input libraries were extracted using the Plasmid Mega Kit (D6926; Omega) and quantified with a NanoPhotometer (Westburg; Leusden, Netherlands).

Illumina HiSeq-compatible primers were used for sequencing. For each biological replicate, the index primers were used in independent final PCR reactions. Each index PCR was performed in 15 replicate PCR reactions (Supplemental Table S8). The PCR products were size-selected (450–500 bp) using a 1% agarose gel (D4008; ZYMO) and submitted for Illumina sequencing.

### Construction and infiltration of *Agrobacterium* input libraries

The plasmid input libraries were introduced into *Agrobacterium* tumefaciens strain GV3101 carrying the helper plasmid pSoup via electroporation. The transformed *Agrobacterium* culture was grown in 150 mL of YEP medium (1% [w/v] yeast extract, 2% [w/v] peptone) until the OD reached 1.0. The bacteria were harvested and resuspended in infiltration buffer (10 mM MES, pH 5.2; 10 mM MgCl2; 0.5 M MES, pH 5.6; and 100 mM AS) to an OD of 1. *N. benthamiana* was grown in soil at 25°C in a long-day photoperiod (16 h light and 8 h dark; cool-white fluorescent lights [Philips TL-D 58W/840]; intensity 300 μmol/m2/s). The suspension was then infiltrated into the two oldest fully expanded leaves of 20 vigorously growing *N. benthamiana* plants (approximately one month old, pre-flowering). The plants were maintained under normal growth conditions for 72 hours.

Illumina HiSeq-compatible primers were used for sequencing. For each biological replicate, the index primers were used in independent final PCR reactions. Each index PCR was performed in 15 replicate PCR reactions (Supplemental Table S8). The PCR products were size-selected (450–500 bp) using a 1% agarose gel (D4008; ZYMO) and submitted for Illumina sequencing.

### Preparation of Agro-STARR-seq output Libraries

A total of 2000 μg of total RNA was extracted from the transformed *N. benthamiana* leaves using the Plant RNAexr Kit (02160037; HYY). Polyadenylated (poly A) mRNA was then isolated using the Dynabeads® Oligo(dT) kit (61005; ThermoFisher Scientific). The mRNA was treated with TURBO™ DNase (AM2239; ThermoFisher Scientific) and purified using Agencourt RNAClean XP beads (A66514; Beckman Coulter).

First-strand cDNA was synthesized using SuperScript III (18080093; ThermoFisher Scientific) with 1.5 μg mRNA per reaction. Target-specific reverse transcription (RT; Supplemental Table S8) was performed using Agro-STARR-seq mRNA-specific RT primers. All mRNA samples were used for the RT reaction. After RT, the reaction was treated with RNase A and H, followed by pooling. In the final PCR amplification, 5 μL of cDNA reaction was used as a template for a 50 μL PCR reaction with the KAPA HiFi HotStart ReadyMix PCR Kit (KK2602; Roche). The PCR cycling conditions were: 98°C for 30 s; 20 cycles of 98°C for 45 s; 15 cycles of 98°C for 15 s, 65°C for 30 s, and 72°C for 70 s.

Illumina HiSeq-compatible primers were used for sequencing. For each biological replicate, 24 different index primers were used in independent final PCR reactions. Each index PCR was performed in 15 replicate PCR reactions (Supplemental Table S8). The PCR products were size-selected (450–500 bp) using a 1% agarose gel (D4008; ZYMO) and submitted for Illumina sequencing.

### Luciferase Reporter Assay

For the luciferase reporter assay, the vector backbone used for the reporter and control plasmids was identical to that of the plasmid input library but lacked the CmR-ccdB cassette (Fig. S8A and S8B). The candidate regions for the reporter plasmids were derived from *Arabidopsis* genomic DNA. A dual-luciferase assay was performed in *N. benthamiana* with three independent biological replicates. The dual-luciferase reporter plasmid was introduced into *Agrobacterium* strain GV3101 via chemical transformation for transient expression in *N. benthamiana* leaves. After 72 hours, the dual-luciferase activity was measured following the manufacturer’s protocol (E1910; Promega).

### RNA isolation, RT-PCR, and qRT-PCR

RNA was extracted from the *N. benthamiana* leaves transformed with *Agrobacterium*. To identify EGFP genes, the suspension was then infiltrated into the two oldest fully expanded leaves of 20 vigorously growing *N. benthamiana* plants (approximately one month old, pre-flowering). The plants were maintained under normal growth conditions for 72 hours. and subjected to RNA extraction with RNAiso Plus (9109; Takara). First-strand cDNA was generated via reverse transcription reaction with a HiScript ®II 1st Strand cDNA Synthesis kit (R312-01; Vazyme). The transcript levels of LORELEI family genes, ROPs and NADPH oxidases were measured by RT-PCR with the primer pairs listed (Supplemental Table S8). The abundance of RALF transcripts was detected using qRT-PCR with ChamQ Universal SYBR qPCR Master Mix (Q711-02; Vazyme). The following program was used: 95℃ for 5 min, followed by 30 cycles at 95℃ for 10 s, and 60℃ for 30 s. The relative transcript abundance was calculated using the 2–ΔΔCT method. The numbers were presented as means ± SD from three replicates.

### Bioinformatic Analysis of Agro-STARR-seq in *Arabidopsis*

#### Agro-STARR-seq Data Analysis

Agro-STARR-seq involves two datasets with different types of readouts. The input libraries were generated by directly amplifying the inserted fragments from the plasmid DNA used for *Agrobacterium* electroporation, representing the original composition of the starting *Agrobacterium* plasmid library. The output libraries measure the abundance of self-transcribed mRNA from the inserted fragments in the transfected *Agrobacterium* plasmid library.

### Enhancer peak calling and Distribution

Fastp (v0.23.4) was used to filter low-quality reads from raw sequencing data with the parameter “-w 5”, while all other parameters remained at their default settings. The filtered reads were then aligned to the TAIR10 reference genome using Bowtie2 (v2.2.8) with the following settings: “--wrapper basic-0 --very-sensitive -X 2000 --rg-id <index id>”. Duplicate reads from different index libraries were removed separately using Sambamba. Enhancer peak calling was performed with MACS2 (v2.1.4) using the command: “callpeak -c <STARR-seq input library bam> -t <STARR-seq output library bam> --keep-dup all -q 0.05 --gsize 119667750 –nomodel”. R package "ideogram" was used to plot the genome-wide distribution of enhancers.

### Enhancer annotation and conservation analysis

The distribution of enhancers was analyzed using the function ‘annotatePeak’ from R package ChIPSeeker [24] (v1.42.1) with genomic annotation provided by R package ‘TxDb.Athaliana.BioMart.plantsmart51’.

PhastCons conservation scores were obtained from PlantRegMap [25], and genomic region annotations were downloaded from Ensembl Plants. The average PhastCons score was calculated for each enhancer and genomic region for comparative analysis.

### Epigenetic states analysis

Epigenomic datasets for *Arabidopsis thaliana* were obtained from published studies in the GEO database: GSE183957 (mature leaves, Supplemental Table S9) [26] and GSE233525 (young shoots, Supplemental Table S9) [16]. These datasets include H3K27ac, H3K9ac, H3K27me3, H3K4me1, H3K4me3, and open chromatin. Data processing, including trimming, mapping, and filtering, was performed for subsequent analysis.

### Motif analysis

A total of 619 transcription factors (TFs) of *Arabidopsis thaliana* were downloaded from PlantTFDB v5.0 [25]. Motif enrichment analysis was performed using a custom script. The background for motif enrichment analysis was set as shuffled regions of non-enhancer regions. ERF115 was selected as an TF example and visualized in IGV (Fig. S4E, S4F and S4G), based on data obtained from the published study GSE48793 (Supplemental Table S9) [27].

### Transposable elements

Annotated transposable elements (TEs) were downloaded from the *Arabidopsis* Information Resource (TAIR, TAIR10 genome release).

### Deep learning model training and enhancer design (DREAM framework)

Dataset Composition: To enhance data diversity while ensuring accurate quantification of enhancer activity, we randomly selected background genomic regions equal in number to the peak regions, with input DNA read counts > 30. To capture comprehensive enhancer sequence information, we extended 300 bp in both directions from the midpoint of each enhancer peak. Enhancer activity was quantified as the log2 fold change of RNA reads RPKM mapped to the genomic region over the input DNA reads RPKM. The dataset was further augmented by adding the reverse complement of each sequence with the same regulatory activity. Genomic regions on chromosomes 4 and 5 were used as validation and test datasets respectively, while the remaining regions served as training data. Model training and enhancer design: We employed our previously published DREAM framework for training, with a learning rate of 4e-5 and batch size of 256, while other parameters remained consistent with our prior research [24]. In the genetic algorithm, the initial population size, mutation rate, and other parameters were maintained as in our previous work [14].

### Bioinformatic Pipeline and Analysis of ATAC-Agro-STARR-seq in *N. benthamiana*

We generally followed the ATAC-STARR-seq pipeline as described by Hansen TJ. et al. 2022 [17].

### Read processing

Reads from ATAC-seq, plasmid, and Agrobacterium-mediated input (DNA) and output (RNA) libraries were processed using fastq_processing.py. The reference genome and gene annotation files were based on Niben261 (downloaded from http://solgenomics.net/), with chloroplast (C_AA066594.1) and mitochondrial (C_AA066595.1) sequences obtained from CNCB. For enhancer identification, duplicate reads were retained in the *Agrobacterium*-mediated DNA and RNA libraries, as suggested in the first ATAC-STARR-Seq paper [17]. The summary statistics are provided in Supplemental Table 3. These duplicate-retained reads were also used to estimate library complexity with the lc_extrap function from Preseq (https://github.com/smithlabcode/preseq, version 3.2.0) and to analyze insert size distribution with the CollectInsertSizeMetrics tool from Picard (https://broadinstitute.github.io/picard/, version 2.26.3).

### Accessibility Analysis

Genrich (https://github.com/jsh58/Genrich, version 0.6.1) was used to call accessibility peaks for ATAC-seq. The gene distribution of ATAC-seq peaks and enhancer regions as well as active histone modification peaks (described below) was analyzed using the ChIPSeeker package. The promoter region (“tssRegion” parameter) was defined as 2000 bp upstream and 1000 bp downstream of the TSS. The *Niben261* TxDb object was constructed using the “makeTxDbFromGFF” function from the GenomicFeatures R package [28].

For TSS enrichment analysis, gene locations and strand orientations were extracted from the GFF annotation file. The “computeMatrix reference-point” and “plotProfile” functions from deepTools (version 3.5.6) were then used to generate TSS enrichment profiles.

To compare the expression levels of genes associated with DNA-accessible proximal and distal regions, we downloaded three replicate *N. benthamiana* LAB leaf RNA-seq datasets (SRR25703579, SRR25703580, and SRR25703581) (Supplemental Table S9) [29]. Low-quality reads were filtered using Trim Galore (https://github.com/FelixKrueger/TrimGalore), and the remaining reads were aligned to the reference genome with HISAT2 (version 2.2.1) [30]. Reads with a mapping quality score (MAPQ) ≥ 30 were retained for expression quantification. Gene-level read counts were obtained using HTSeq-count (version 0.11.3) [31], and expression values were normalized as FPKM using the fpkm function in edgeR [32]. The mean FPKM across the three replicates was calculated for each gene.

### Enhancer Identification and Characterization

Duplicate-retained DNA and RNA libraries were used as input for the call_ATAC-STARR_regulatory-regions.py script to identify active bins and active regions, with ATAC-seq peaks provided as the input peak set. Active regions were classified as proximal or distal using the “annotatePeak” function from the ChIPSeeker package. Active regions located within promoter regions were considered proximal active regions, while those outside this region were considered distal active regions.

For histone modification analysis in active regions, we downloaded *Nicotiana benthamiana* histone modification ChIP-seq data [29], including SRR27031034 (H3K27ac), SRR27031032 (H3K4me3), SRR27031033 (H3K27me3), SRR27031030 (H3K9me2), and their input SRR27031035, SRR27031031 (Supplemental Table S9). And we used the “computeMatrix reference-point” and “plotHeatmap” functions from deepTools to generate heatmaps for proximal and distal regulatory regions.

To compute transcription factor (TF) footprints within accessible peaks, we applied the TOBIAS software to the DNA library. The analysis followed a similar approach to Hansen et al. [17], except that *Arabidopsis thaliana* TF binding motifs were used, obtained from PlantTFDB. Bound and unbound transcription factors (TFs) were defined using the BINDetect step. TCP ChIP-seq datasets were obtained from SRR15036661 and SRR15036676.

For motif enrichment analysis, TF motifs from *Arabidopsis thaliana* were first scanned in all active regions, proximal active regions, distal active regions, and background regions using FIMO [33]. Background sequences consisted of 50,000 randomly selected regions from the reference genome, excluding active regions, with an average length of 115 bp, corresponding to the mean active region length. Then, the number of regions containing each motif was counted for each type of region. Finally, we used a chi-square test to identify motifs that were significantly enriched compared to the background.

To analyze the functions of genes associated with TCP-bound active accessible peaks, we extracted gene sets annotated to the bound peaks for each TCP transcription factor. We performed GO enrichment analysis for these gene sets using clusterProfiler [34]. GO term annotations were obtained from the eggNOG-mapper website [35] (http://eggnog-mapper.embl.de/).

Because no CCA1-like gene is annotated in Niben261, we extracted RNA-seq reads from the region surrounding Niben261Chr12:120,946,470–120,948,272 and used StringTie [36] (version v3.0.0) to predict transcripts. The resulting exon sequences were then aligned to *Arabidopsis* Araport11 coding sequences using blastn in website: (https://www.arabidopsis.org/) to confirm its homologous gene.

## Acknowledgements

We would like to thank Yang Fu, Hu Yang, Yantao Wang, Renyu Deng, Sheng Yu and Zhenghang Zhu for their technical assistance.

## Author’s contributions

Y.L. conceived the experiments. Y.B. conducted the experiments. H.L., J.Q., H.C., N.L., Y.C., L.X., Y.Z., F.Z and C.C. provided the technical support. Z.L. and X.Z. led the bioinformatics analysis; Z.L., X.Z., Y.S., and W.Z. performed the bioinformatics analysis; S.Z. supervised the work; Y.B., X.Z., Z.L. and Y.L. wrote the manuscript, and all authors approved the final manuscript.

Y.B., X.Z. and Y.S. should be considered co-first authors.

## Data availability

Raw sequencing data will be made available upon publication.

## Declarations

### Ethics approval and consent to participate

Not applicable.

